# Ligands with poly-fluorophenyl moieties promote a local structural rearrangement in the Spinach2 and Broccoli aptamers that increases ligand affinities

**DOI:** 10.1101/2021.10.05.463302

**Authors:** Sharif Anisuzzaman, Ivan M Geraskin, Muslum Ilgu, Lee Bendickson, George A Kraus, Marit Nilsen-Hamilton

## Abstract

The interaction of nucleic acids with their molecular targets often involves structural reorganization that may traverse a complex folding landscape. With the more recent recognition that many RNAs, both coding and noncoding, may regulate cellular activities by interacting with target molecules, it becomes increasingly important to understand the means by which nucleic acids interact with their targets and how drugs might be developed that can influence critical folding transitions. We have extensively investigated the interaction of the Spinach2 and Broccoli aptamers with a library of small molecule ligands modified by various extensions from the imido nitrogen of DFHBI (3,5-difluoro-4-hydroxybenzylidene imidazolinone) that reach out from the Spinach2 ligand binding pocket. Studies of the interaction of these compounds with the aptamers revealed that poly-fluorophenyl-modified ligands initiate a slow change in aptamer affinity that takes an extended time (half-life of ~40 min) to achieve. The change in affinity appears to involve an initial disruption of the entrance to the ligand binding pocket followed by a gradual lockdown for which the most likely driving force is an interaction of the gateway adenine with a nearby 2’OH group. These results suggest that poly-fluorophenyl modifications might increase the ability of small molecule drugs to disrupt local structure and promote RNA remodeling.

## INTRODUCTION

Ligand-induced changes in RNA structure and function are important aspects of cellular regulation and their understanding is critical for the development of drugs that alter RNA function and structure *in vivo*. The interactions of RNAs with their target molecules frequently involve significant changes in RNA structure, even if only local. These local changes are amplified in riboswitches and other functional RNAs with resulting changes in local and global RNA structure (Mandal et al. 2004; Suess et al. 2004; Priyakumar 2010; Stoddard et al. 2010; Dethoff et al. 2012).

The paths by which structural changes occur in RNAs have not frequently been studied at the molecular level in part due to the difficulty of tracking such changes over time. Spinach and Broccoli are light-up aptamers that provide an environment conducive for an increase in ligand fluorescence when bound. As this feature is compatible with tracking aptamer binding capability over long time periods we have investigated the interactions of a series of derivatives of the ligand, DFHBI (3,5-difluoro-4-hydroxybenzylidene imidazolinone), with the Spinach2 and Broccoli aptamers to evaluate the stability of these molecular structures over time.

Here we describe a new interaction of two poly-fluorophenyl ligands with the Spinach2 and Broccoli aptamers that involves a local structural rearrangement reflected in a slow increase in ligand affinity and requires the presence of multiple fluorines on a benzene ring appended to the imido nitrogen of DFHBI.

The rearrangement results in a remodeled local structure that involves a predicted single H bond between the amino group of the gateway adenine and a close-by 2’OH group. This transition demonstrates how a small ligand can alter the local structure of a much larger, already folded, RNA. The pathway in Spinach2 appears to be an initial disruption of the local RNA structure followed by relaxation to an alternative structure in which the binding pocket is locked down by a hydrogen bond between the gateway adenine and a nearby 2’ hydroxyl group. Our results suggest that drugs directed to functional RNAs are likely to be more effective in altering RNA functionality if they contained moieties such as multiply fluorinated phenyl groups with the ability to disrupt local RNA structures and promote rearrangement to new ligand-driven structures.

## RESULTS

### Ligand screen of Spinach and Broccoli aptamer affinities

Studies of the binding affinities of the Spinach aptamer family (Spinach, Spinach2, Baby Spinach, iSpinach, and Broccoli) have consistently identified an adenosine group, referred to as the “gateway adenine” or “lid layer”, as important for ligand affinity (Huang et al. 2014; Warner et al. 2014; Ageely et al. 2016; Warner et al. 2017). This adenine is present in position 69 in the deposited crystal structure of Spinach2 (PDBID;4TS2) and in position 71 in SPN2A. The gateway A is seen in the crystal structure very close to the imido-amino-methyl group of DFHBI, which protrudes from the Spinach2 aptamer binding pocket (Fig. 1A) and is shown to make a direct contact with the methyl group of DFHBI in the crystal structure of iSpinach (Fernandez-Millan et al. 2017).

**Figure 1:**
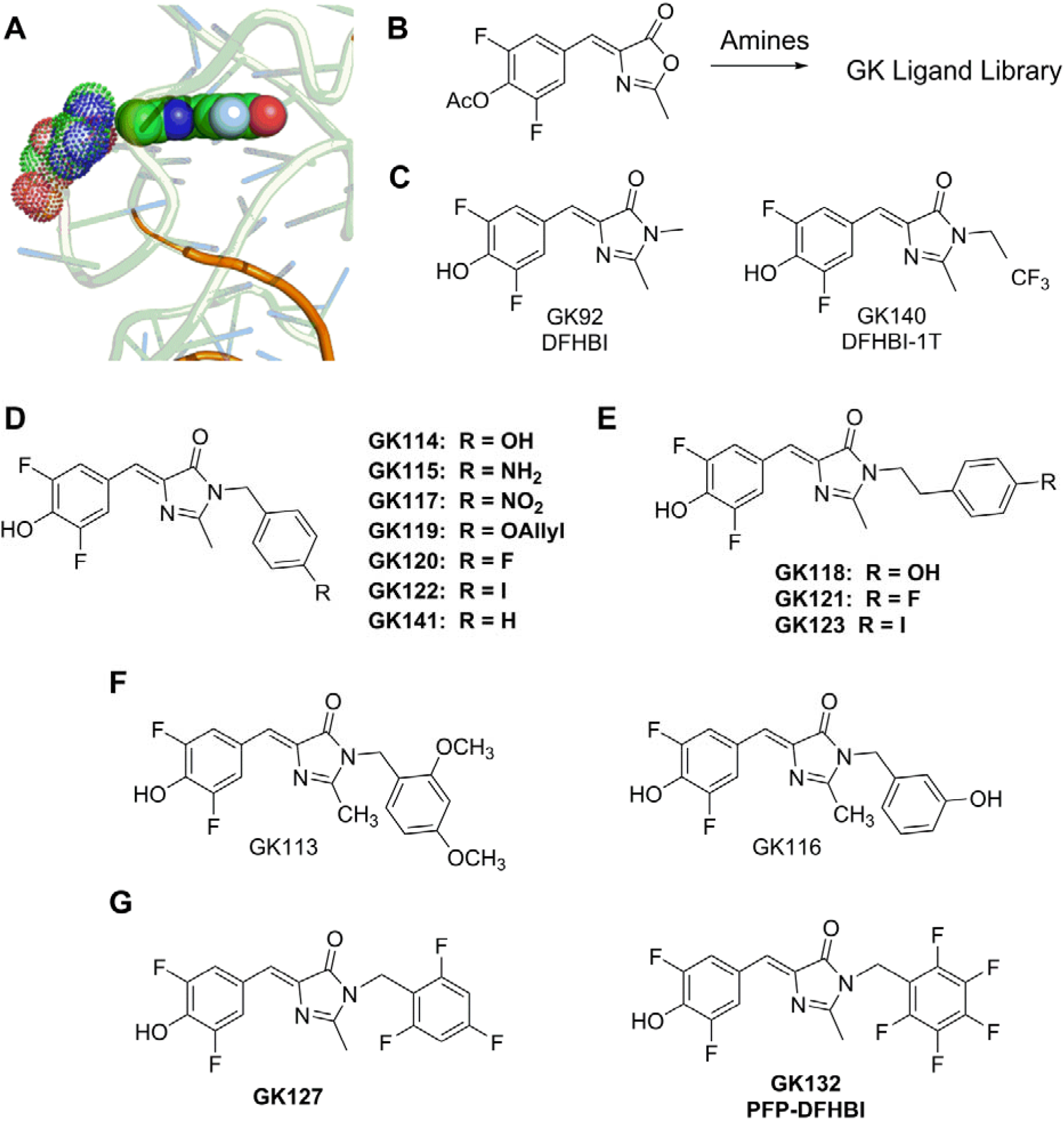
DFHBI imido-N extension library. **A)** Positions of the gateway A (stippled) and the DFHBI methyl group (filled) Spinach2 crystal structure. **B)** Overview of the synthesis protocol for GK compounds **C-G)** Chemical structures of the synthesized DFHBI derivatives.

To better understand the means by which the gateway A influences ligand affinity, a library of DFHBI derivatives was produced in which the amino methyl group was extended with a variety of phenyl and benzyl groups (synthetic outcomes in Supplemental materials). Selection of the Spinach aptamer was performed with DFHBI linked to a solid support by way of the imido nitrogen (Song et al. 2014). We have found this position to be optimal for modifying DFHBI while maintaining aptamer affinity for the newly-synthesized modified ligands. To create the library of alternative DFHBI ligands, azlactone (Fig. 1B) was reacted with primary benzylic amines and phenethylamines (Fig. 1C). By this means, DFHBI was modified by a variety of phenyl or benzyl groups linked to its imidazole nitrogen (Fig. 1D–G). Of these compounds GK92 (DFHBI) and GK132 (PFP-DFHBI) have been previously reported (Paige et al. 2011; Ilgu et al. 2016). It was hypothesized that the phenyl or benzyl extensions from DFHBI might interact with the gateway A to either increase affinity by stacking or to decrease affinity by preventing the A from forming a critical interaction.

The photophysics of DFHBI and PFP-DFHBI in the absence of aptamer were examined in detail and both were found to have similar photoisomerization characteristics and pH dependence of fluorescence (Santra et al. 2019). We determined the quantum yield for DFHBI in complex with tandem-SPN2A as 0.71, which compares favorably with previously reported value of 0.72 (Paige et al. 2011). The quantum yield for PFP-DFHBI in complex with tandem-SPN2A was 0.85 ± 0.053 (N=2).

The Spinach2 (SPN2A) and Broccoli (BRC1A) aptamers were tested for their affinities with various members of the ligand library (Fig 2). Although addition of the phenyl group (GK141) had a small impact on SPN2A affinity, additional constituents to the phenyl ring such as amino (GK115), O-allyl (GK119) and fluorine had large effects resulting in over a 20-fold range of affinities for SPN2A, depending on the constituents (Fig. 2A, Table S1).

**Figure 2:**
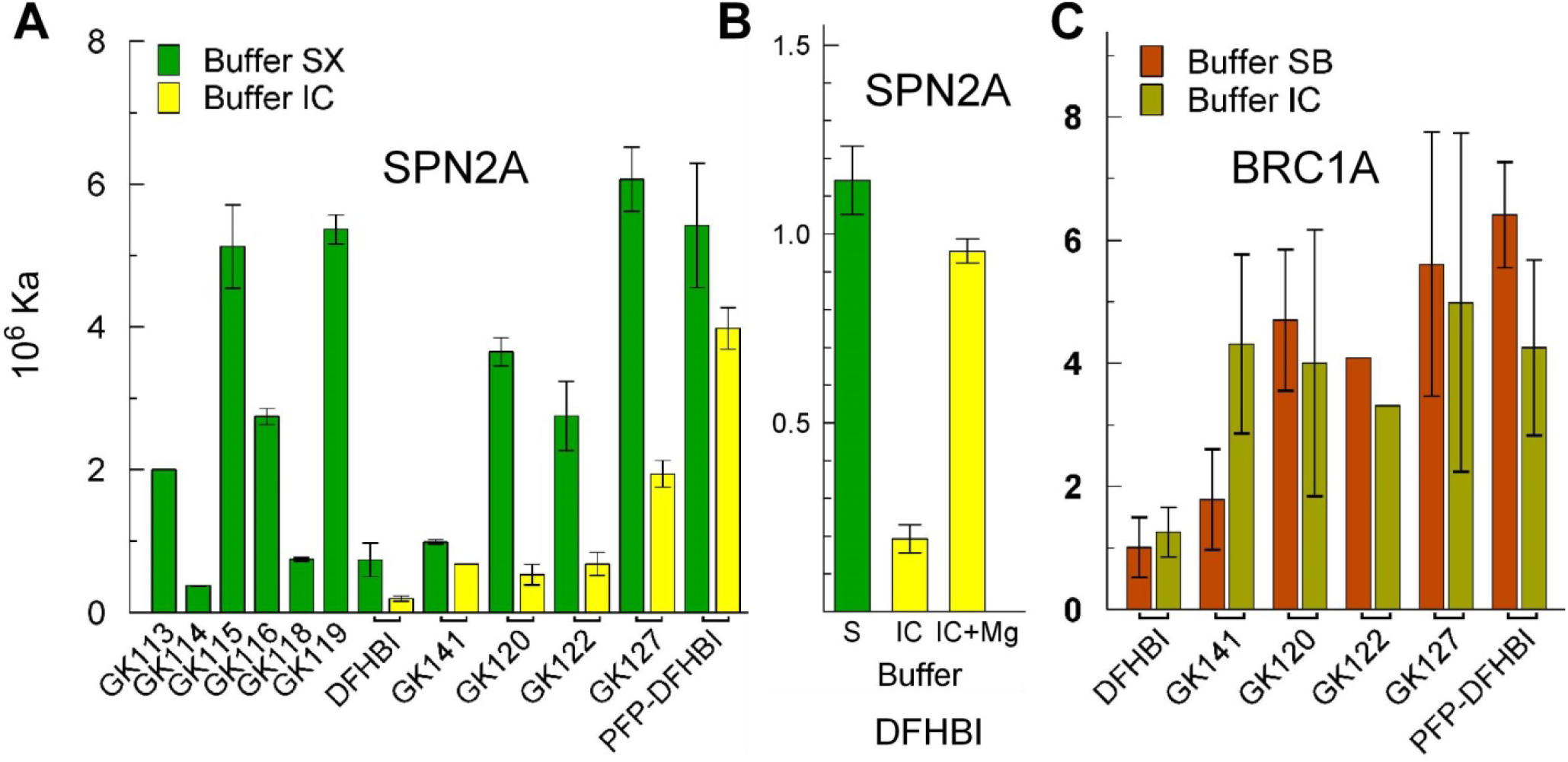
Affinities of SPN2A and BRC1A for imido-N extension ligands. **A)** The affinities of SPN2A for compounds of the imido-N extension ligand library in buffer SX (green bars) or buffer IC (yellow bars) after 10 min incubation. **B)** The effect of Mg on the affinity of SPN2A for DFHBI. Buffer IC (green bar), Buffer IC made 5 mM Mg (yellow bars). **C)** The affinities of BRC1A for some of the imido-N extension ligands in buffer SB (brown bars) or buffer IC (tan bars). All affinities were determined by fluorescence spectroscopy as described in Materials and Methods.

The original Spinach aptamer was selected in buffer SX, which contains 5 mM MgCl_2_ (Paige et al. 2011). Reports of the interaction of its derivative aptamer, Spinach2, with potential ligands were performed using the same buffer but with 1 mM MgCl_2_ (Strack et al. 2013). The binding studies shown in figure 2A and B (green bars) were performed in buffer SX. However, in the mammalian cell, in which this aptamer has been applied, the free MgCl_2_ concentration is about 220 μM (Romani 2013). Therefore, we also tested the binding affinity of SPN2A for the DFHBI imido-N extension library in buffer IC, formulated to resemble the salt content and pH of the mammalian cytoplasm. The results (Fig. 2A, yellow bars) show that DFHBI binds with lower affinity to SPN2A in IC buffer than in SX buffer and this difference between buffers decreases with the addition of three or five fluorines on the phenyl ring. The critical difference between Buffers SX and IC for SPN2A ligand affinity is the MgCl_2_ concentration (Fig. 2B).

The Broccoli aptamer was selected from a Spinach aptamer-derived library in a buffer with a lower concentration of MgCl_2_ to overcome the requirement for high Mg^2+^ (Filonov et al. 2014). Broccoli bound to DFHBI and derivatives equally well in both Buffers SB and IC with the possible exception of GK141 (Fig 2C). Thus, for both Spinach and Broccoli, the affinity for DHBI is enhanced by the inclusion of a phenyl extension from the imido N-methyl group and is impacted by the constituent(s) on the phenyl ring.

### Location of the phenyl extensions to DFHBI relative to the binding pocket

The addition of a phenyl group to DFHBI increased SPN2A ligand affinity in almost every instance (Fig. 2A). Therefore we postulated that this group might interact with chemical groups on the aptamer near the opening of the binding pocket. If so, then the position of the phenyl group immediately outside the entry to the pocket might be important for affinity. To test this hypothesis, we prepared a series of DFHBI derivatives with phenethyl extensions (GK118, 121, and 123) and compared these with their benzyl extension counterparts (GK114, 120, 122) for their abilities to bind SPN2A (Fig. 3A). As well as extending further from the imido nitrogen than the benzyl extensions, the phenethyl extensions were predicted to take an alternate orientation to the imidazole group when free in solution. The predominant structures of the phenethyl extensions were predicted to be in plane with the imidazole group whereas the benzyl extensions are predicted to orient in a plane almost 90 degrees to the imidazole group (Fig. 3A). In all instances, DFHBI analogs with phenethyl extensions from the imido nitrogen had lower affinity for SPN2A than the equivalent benzyl extensions regardless of the substitution on the phenyl group (Fig. 3B). In two of the three instances, SPN2A bound the analogs with the two-carbon linkers to the imido nitrogen with lower affinity than for DFHBI, which has a methyl group at this position. These results are consistent with the hypothesis that the phenyl extension to DFHBI provides an opportunity for interaction of the phenyl group and its appendages with portions of the aptamer close to the binding pocket.

**Figure 3:**
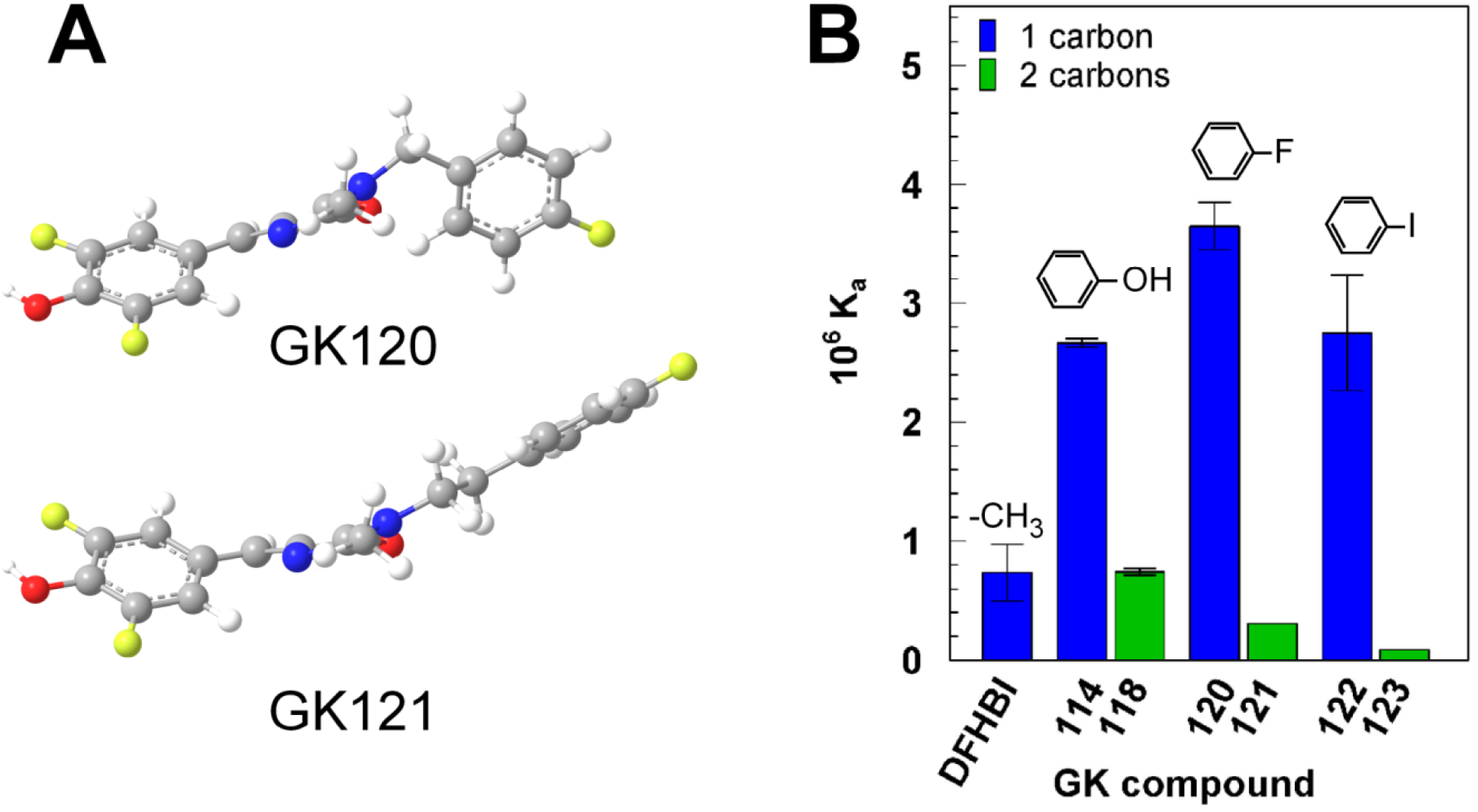
Influence of the position of the phenyl group relative to the imido nitrogen of DFHBI. **A)** The predicted 3D solution structures of two DFHBI derivatives with either a one carbon link to the phenyl group (GK120) or a two carbon link (GK121). **B)** Affinities (K_a_s) in Buffer SX of compounds with benzyl or phenethyl extensions from the imido nitrogen.

### Time-dependent change in Spinach2 and Broccoli aptamer affinity

An unusual feature of the interaction of SPN2A and BRC1A with the DFHBI analogs containing tri- or penta-fluoro substituents was a slow increase of 2 to 3-fold in the affinities of the aptamers for ligand over a period of hours at room temperature (22-24°C; Fig 4A,B). This affinity change was unique for these two ligands and the half-life for the change was 40 ± 4.1 min (N=8). The change in affinity was observed for SPN2A in IC and SX buffer and in SX buffer at pH 7.4 and 8. It was also observed for BRC1A in SB buffer. Although the initial affinities of SPN2A and BRC1A were lower for DFHBI than for PFP-DFHBI, the ability to drive a change in affinity with time was not related to the initial affinity. For example, GK115 and GK119 bind Spinach with similar affinities to GK127 and PFP-DFHBI (Fig. 2A), but only the latter two ligands drive the slow change in affinity with time (Fig. 4A). To ensure that PFP-DFHBI was not changing structurally over this long incubation time, we determined its MS and UV-Vis spectra before and after repeated exposure to 485 nm over the 18 h time period. There was no change in the UV-Vis spectra over this time period (Fig. S3). Mass spectrometry analysis showed the expected peaks after 10 min incubation with one 30s exposure to 485 nm light as were found after thirteen 30 sec exposures to 485 nm over a period of 18 h incubation (Fig. S3 legend). The effect of incubation time on fluorescence in the presence and absence of SPN2A was determined for DFHBI, PFP-DFHBI and DFHBI derivatives. Fluorescence was stable for all compounds in the absence of SPN2A but increased about 20% in the presence of aptamer for most compounds (Table S3). The increase in fluorescence observed with DFHBI and PFP-DFHI was identical and therefore did not contribute to the difference in measured K_a_ for these compounds.

**Figure 4:**
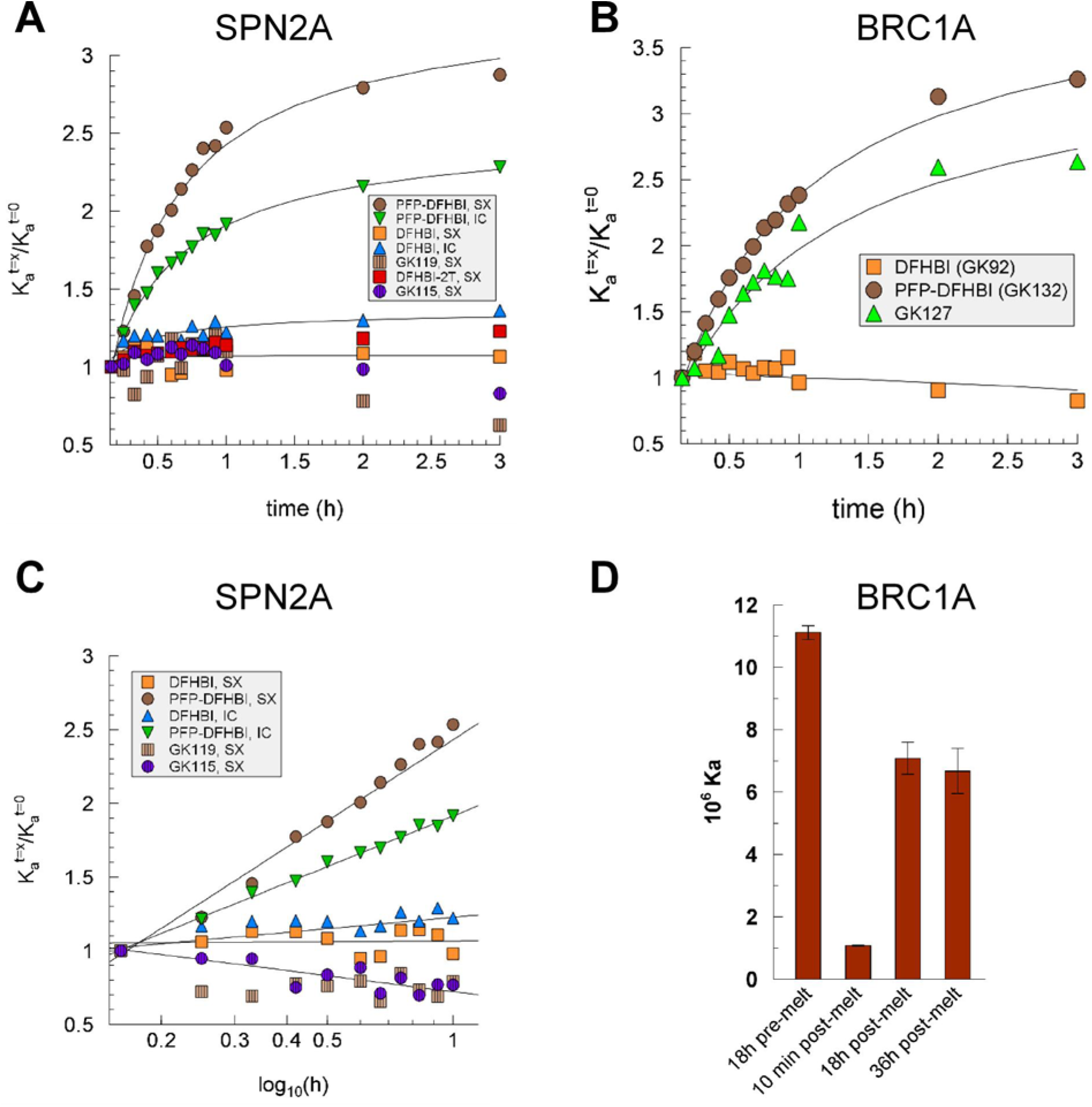
Increase in affinity with incubation time for ligands with polyfluorophenyl imido-N extensions. Affinity as a function of incubation time in the dark for **A)** SPN2A with a variety of DFHBI derivatives in buffers identified in the legend and **B)** Broccoli in Buffer SB. **C)** Data from (A) represented as a semi-log plot. **D)** BRC1A was incubated for 18 h with PFP-DFHBI in Buffer SB (18h pre-melt) then melted at 80°C for 5 min, shifted to 23°C and the affinity measured after 10 min, 18 h and 36 h post-melt.

To further establish the change in affinity over time with PFP-DFHBI and not DFHBI, we determined the on and off rates. These values (Table 1) confirmed that incubation of SPN2A with PFP-DFHBI but not with DFHBI resulted in a change in the kinetics of binding to the respective ligand. The calculated values of K_a_ for both ligands from the k_on_ and k_off_ measurements after 10 min incubation matched well with the K_a_s measured by fluorescence intensity. The k_off_ was measured using GK117 to replace the exited ligand. The ratios of k_off_ (10 min) over k_off_ (18 h) were 1.2 (DFHBI) and 0.72 (PFP-DFHBI). When analyzed for statistical significance using a paired Student’s t-test, the resulting p values were higher than 0.05. Unlike the k_off_, which was not altered for either ligand with longer incubation nor significantly different between ligands, the k_on_ for the two ligands suggested the mechanism by which the ligand affinities are differentiated and how the SPNA affinity for PFP-DFHBI increases with time of incubation. Whereas, the calculated k_on_ for DFHBI at 18 h was 0.84 of the measured value at 10 min, it was 1.66 times the k_on_ measured at 10 min for PFP-DFHBI. This 66% change is well outside the error range of all measurements of K_a_, k_off_ and k_on_ (17% ± 8%). Thus, we conclude that difference in affinities of SPN2A for DFHBI and PFP-DFHBI and the increase in affinity of SPN2A with PFP-DFHBI with incubation is defined by the k_on_.

**Table 1:** Kinetic measurements for the Spinach aptamer interaction with DFHBI and PFP-DFHBI.

| Ligand | DFHBI |  | PFP-DFHBI |  |
| --- | --- | --- | --- | --- |
| Incubation time | 10 min | 18 h | 10 min | 18 h |
| $K_a$ ($\mu\text{M}^{-1}$) | $0.33 \pm 0.04$ (2) | $0.28 \pm 0.30$ (2) | $4.0 \pm 0.84$ (5) | $8.3 \pm 1.0$ (4) |
| $k_{\text{off}}$ ($\text{s}^{-1}$) | $4.9 \pm 1.2 \times 10^{-4}$ (6) | $5.8 \pm 1.3 \times 10^{-4}$ (6) | $5.3 \pm 0.15 \times 10^{-4}$ (2) | $3.9 \pm 0.15 \times 10^{-4}$ (3) |
| $k_{\text{on}}$ ($\mu\text{M}^{-1}\text{s}^{-1}$) | $1.9 \pm 0.48 \times 10^{-4}$ (6) | $1.3 \times 10^{-4}$ (calculated) | $1.9 \pm 0.17 \times 10^{-3}$ (3) | $3.2 \times 10^{-3}$ (calculated) |
| $K_a$ (calculated) | 0.39 | | 3.6 | |

**Legend:** $k_{\text{on}}$ after 10 min and $K_a$ and $k_{\text{off}}$ after 10 min and 18 h incubation were determined for DFHBI and PFP-DFHBI. The $k_{\text{on}}$ at 18 h was calculated from the measured $K_a$ and $k_{\text{off}}$ . The $K_a$ at 10 min was also calculated from the measured $k_{\text{on}}$ and $k_{\text{off}}$ and found to be within 20% of the measured values for both compounds. Shown are the measured values ( ± SEM) with the number of independent estimates in parentheses. For $k_{\text{on}}$ , each independent estimate involved measuring $k_{\text{obs}}$ for at least four different ligand concentrations. For $k_{\text{off}}$ , each independent measurement involved a duplicate measurement of the $k_{\text{off}}$ for the same combination of ligand and aptamer with the duplicates independently constituted and analyzed for $k_{\text{off}}$ on the same day.

The change in affinities observed with GK127 and PFP-DFHBI occurred with the characteristics of first order reactions. This suggested a single event is responsible for shifting the aptamer into a higher affinity state. As fluorine is the constituent of both active ligands, we considered the possibility of an unusual chemical reaction causing the change in aptamer affinities. Being the most electronegative of elements, fluorine is not likely to be displaced from the ligand. But, fluorine can participate in H-bonds and also alters the excited electronic states of benzene by the “perfluoro effect” (Mondal and Mahapatra 2010). To investigate the possibility that the presence of fluorophenyl groups close to the aptamer might result in a covalent reaction between the aptamer and ligand or within the aptamers that locks them in a higher affinity state, we determined if a previous incubation of ligand and aptamer would increase the initial affinity of the Broccoli aptamer for PFP-DFHBI (Fig. 4D). The results show that the effect of ligand on aptamer affinity does not survive a melting of the aptamer-ligand complex. Although it is possible that a heat-sensitive chemical reaction occurs to increase the affinity of these aptamers to the two ligands with the fluorophenyl appendages, we consider this an unlikely explanation for the effect of the fluorophenyl-DFHBI ligands on aptamer affinities. This conclusion is supported by the observation that DFHBI-1T, with a trifluoro-methyl group linked to the imido nitrogen, does not cause a change in SPN2A affinity with time (Fig. 4A).

### The gateway A secondary amino group is required for increased aptamer affinity

We investigated the role of the gateway A in driving the change in affinity for the fluorophenyl DFHBI derivatives as several studies have identified it as the important for DFHBI binding affinity. First, we removed the base from the gateway position 71 in SPN2A. To create the apurinic SPN2A and other aptamers with alternate bases, we used the split aptamer format (Warner et al. 2014), which placed the gateway A in position 21 of one aptamer half. For replacements with natural bases, the appropriate DNA templates were used for *in vitro* RNA synthesis. Regardless of the moiety that replaced the gateway A, the affinity for ligands (DFHBI and PFP-DFHBI) was reduced (Fig. 5A). As for the full-length aptamer, incubation with the ligand increased the affinity of the split aptamer for PFP-DFHBI but not for DFHBI (Fig. 5B left panel). Without the gateway base, the apurinic SPN2A bound PFP-DFHBI with about 100-fold lower affinity than with the base present and did not change in affinity for PFP-DFHBI with time (Fig. 5B,C).

**Figure 5:**
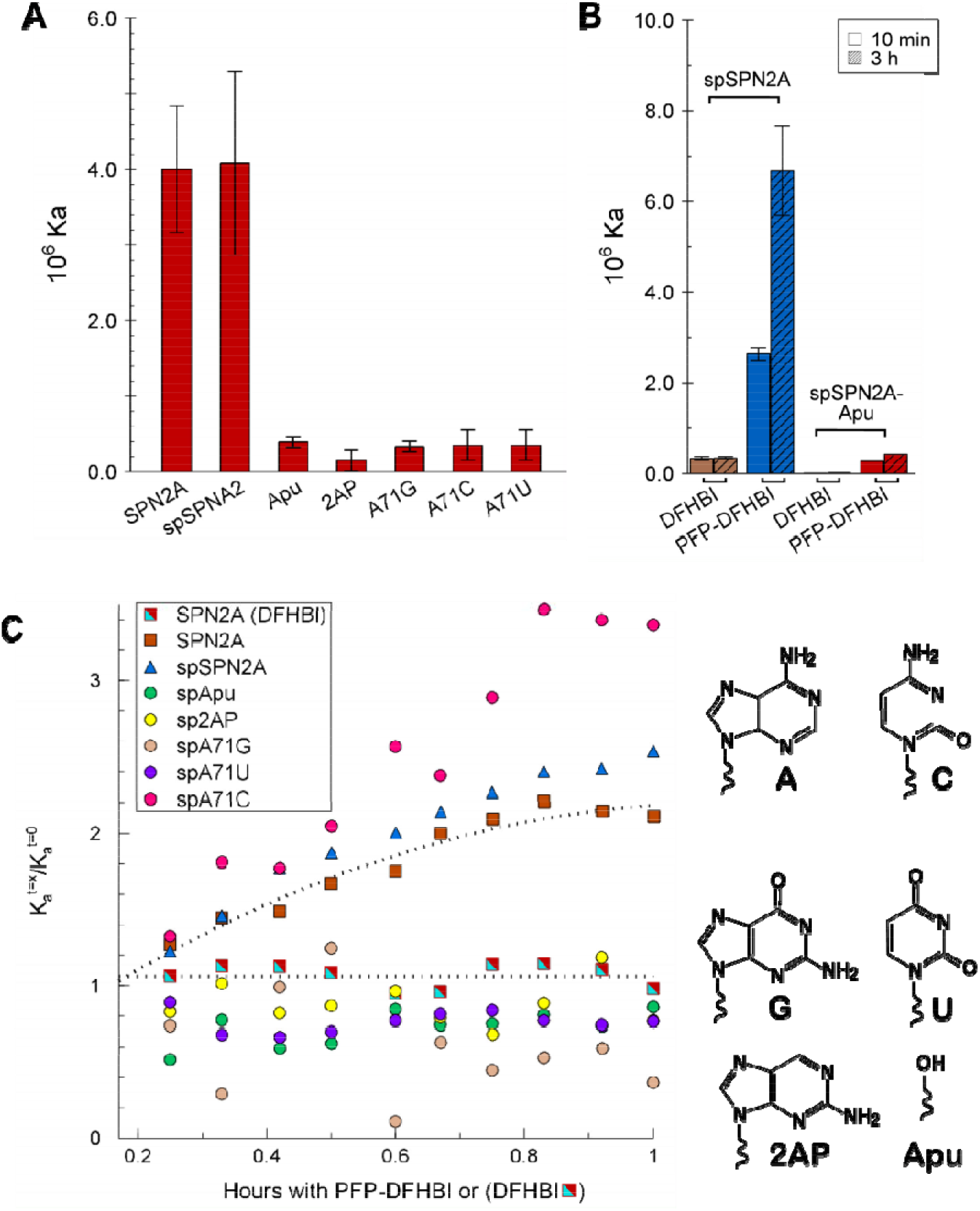
The effects of altering the gateway base on the affinity and time dependent change in affinity of SPN2A for DFHBI and PFP-DFHBI. **A)** Affinities for PFP-DFHBI in Buffer SX of SPN2A variants in position 71 as identified in the X-axis labels. **B)** Comparison of split SPN2A (spSPN2A) and spSPN2A lacking a purine at position 71 for their affinities for DFHBI and PFP-DFHBI after 10 min incubation (open bars) and 3 h incubation at 23°C (hatched bars) in Buffer IC. **C)** Time dependency of the affinities in Buffer SX of SPN2A variants as identified in the legend. All time dependencies are for binding to PFP-DFHBI except the one identified as SPN2A (DFHBI). The glycosidic bond is represented by the squiggle in each chemical structure.

The effect of ligand on aptamer affinity was compared for SPN2A with a deletion or base substitutions for the gateway A (Fig. 5C). The change in affinity with time was observed if A or C were present at the gateway position but not if G, 2AP, U or no base were present. The results suggest that the amino group of A or C might be responsible for forming a hydrogen bond that increases stability of the binding pocket and thereby increases affinity. That SPN2A variants with bases containing a secondary amino group (but not trans to the glycosidic bond) did not increase in affinity with time of incubation suggests that there is a steric component to the hypothesized additional H-bond.

### Local shape changes in Spinach with incubation time and the H-bond partner

To understand the nature of the changes occurring around the ligand binding pocket, SHAPE (Merino et al. 2005; Wilkinson et al. 2005; Steen et al. 2011) was performed with SPN2A in the presence of DFHBI or PFP-DFHBI at 10 min or after 2 h incubation at room temperature. Although the results showed a variety of changes in 2’OH exposure over the entire molecule (Fig. 6A), the most prominent differences between DFHBI and PFP-DFHBI in their interaction with SPN2A over time were around the binding pocket (positions 60-75). After 10 min incubation with either DFHBI or PFP-DFHBI, the region around the gateway A was altered in its exposure to modification by benzoyl cyanide, the SHAPE reagent (Fig. 6B). However, a difference between the effects of these two ligands was observed when 10 min and 2 h incubations were analyzed. Whereas the aptamer shape around the binding pocket changed little with DFHBI over this time period (Fig. 6C), this region appeared destabilized after 10 min with PFP-DFHBI followed by a tightening down after 2 h (Fig. 6D). These results are consistent with the interpretation that PFP-DFHBI disrupts the local RNA structure enabling the aptamer to overcome a local folding energy barrier, which allows formation of a new bond near the ligand binding pocket.

**Figure 6.**
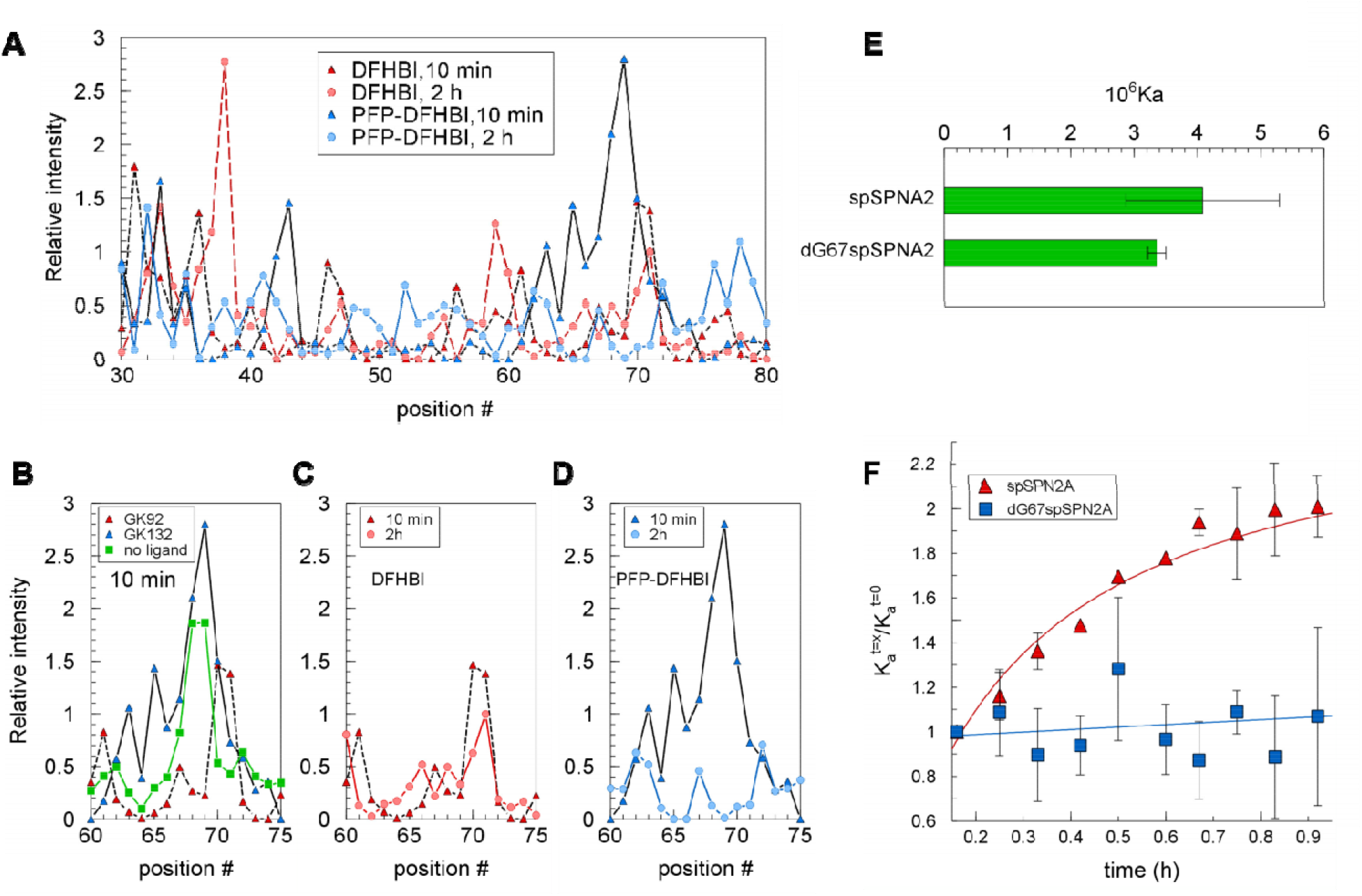
Evidence for an initial disruption by PFP-DFHBI around the SPN2A binding pocket entrance followed by lock-down around the pocket requiring the 2’OH on G67. **A-D)** SPN2A was analyzed by SHAPE after 10 and 120 min incubations with DFHBI and PFP-DFHBI in Buffer SX. **A)** Relative intensities at all aptamer positions from incubations for 10 or 120 min with DFHBI or PFP-DFHBI. **B-D)** Relative intensities in the region around th gateway A after incubation for **B)** 10 min with DFHBI or PFP-DFHBI, **C)** 10 or 120 min with DFHBI or **D)** 10 or 120 min and PFP-DFHBI. **E)** The affinities for PFP-DFHBI of spSPN2A and dG67spSPN2A at 10 min. **F)** Time course of the change in affinity for PFP-DFHBI of SPN2A and SPN2A with dG at position 67 (dG67SPN2A).

We examined the crystal structure of Spinach2 to identify possible candidates for a hypothesized H bond with the secondary amino group of the gateway adenine. The closest hydroxyl to the gateway amino group is on G67 (G65 in the crystal structure), which is 5.5 Å from the amino nitrogen of the gateway A (Fig. S2). This distance is too long to support a hydrogen bond, but we hypothesized that PFP-DFHBI disturbs the aptamer structure around the binding pocket sufficiently to allow a realignment of residues. The result of this disturbance could be the formation of a new H-bond that locks the A over the binding pocket with a resulting increase in ligand affinity. To test this hypothesis, we determined the rate of change in affinity of SPN2A with a dG in position 67 in place of G. Replacement of the deoxyribose for ribose at position 67 did not alter ligand affinity (Fig. 6E). However, unlike the aptamer with a 2’ OH at position 67, the aptamer with 2’ deoxy pentose in position 67 did not change in affinity with time (Fig. 6F). Thus, the 2’ OH at position 67 is required for a change in affinity of SPN2A.

## DISCUSSION

Members of the Spinach aptamer family all possess an adenine in the equivalent gateway position directly outside the ligand pocket entrance (Warner et al. 2014; Fernandez-Millan et al. 2017). The presence of this A is essential for maintaining high ligand affinities of Spinach and related aptamers as shown here and previously by others (Warner et al. 2014; Fernandez-Millan et al. 2017). The crystal structure shows that the gateway A interacts in a T-stack formation with the methyl group of the imidazole nitrogen in DFHBI (Huang et al. 2014). Such an arrangement should readily be replaced by another purine – either 2AP or G. But, any replacement of the A in the gateway position, including of 2AP, resulted in a decrease in affinity. We suspect that the two other purine replacements, 2AP and G, at position 71 might be displaced by interaction(s) with another portion of the aptamer that leaves them unable to replace A as “lid” for the binding pocket.

Here we provide evidence of an alternative structure for Spinach2 with increased ligand affinity, which is achieved as a result of a local rearrangement of the bases around the ligand pocket initiated by poly-fluorophenyl-DFHBI ligands. Our evidence suggests that the new structure is stabilized by a hydrogen bond between the gateway A and the close-by 2’ hydroxyl group of G67. Based on the SHAPE analysis, we speculate that the aptamer rearrangement results from a disturbance at the ligand pocket entrance by the presence of the poly-fluorophenyl moieties of the relevant DFHBI analogs. The direct involvement of the amino group on the A in the rearrangement is supported by the observation that C is the only base other than A that enables an increase in affinity with time. This is despite the observation that the affinity of the aptamer with C71 is very low for PFP-DFHBI.

Two properties of the poly-fluorinated phenyl moieties are possible drivers of local structural disruption. First, unlike other halogens, F can form hydrogen bonds, albeit weak (Howard et al. 1996; Bissantz et al. 2010; Dalvit et al. 2014). Second, fluorines are particularly hydrophobic (Bissantz et al. 2010; Mondal and Mahapatra 2010). Either or both properties might be responsible for a local disturbance in the ligand pocket entrance structure. The possibility of a chemical modification of the aptamer that either involves the ligand or is initiated by the ligand is unlikely. First, the change in affinity due to PFP-DFHBI is completely reversible upon melting the SPN2A structure. Second, the change in affinity is not initiated by interaction with DFHBI-1T, which has a trifluoromethyl extension to the imido nitrogen of DFHBI.

The initial structural disturbance caused by the poly-fluorophenyl appendage to the ligand is proposed to enable sufficient mobility in the region of the pocket entrance for the gateway A amino group and the G67 2’OH to approach each other and form a hydrogen bond. This bond and the microenvironment in which the base is then located is proposed to be sufficiently stable as to behave as a lock on the structure of the pocket entrance. Instead of a direct H-bond between the gateway A and G67, other more complicated interactions may occur that involve these groups and others on the RNA. However, the first order nature of the affinity change with time identifies a single interaction as the driving force for increased aptamer affinity for the poly-fluorophenyl-DFHBI derivatives. Thus, without evidence of the involvement of other components on the RNA, we propose that interaction with PFP-DFHBI results in the formation of a hydrogen bond between A71 and G67, which tightens the pocket entrance and decreases k_off_ with a consequent increase in ligand affinity.

Analysis of the kinetic parameters behind the affinities of Spinach2 for DFHBI and PFP-DFHBI show that the k_off_ is the same for both ligands and does not change with incubation time. Although k_off_ for DFHBI has previously been calculated based on measurements of k_on_ (Han et al. 2013), this is the first time to our knowledge that k_off_ has been measured experimentally. Measurement of k_off_ requires the availability of a high affinity ligand for Spinach2 (GK117) that does not fluoresce when bound to the aptamer. The measured off rates for both ligands after short or long incubation times were very similar and differences did not account for the different affinities of Spinach2 for the two ligands or for PFP-DFHBI after an incubation period. The changes in kinetic parameters that are consistent with the changes in affinity between ligands and incubation time were found in k_on_.

The observed difference between Spinach2 binding of PFP-DFHBI and DFHBI in k_on_ and not in k_off_ suggests that the K_a_ is controlled by the k_on_. A change in k_off_ would be consistent with a change in the number or type of interactions between aptamer and ligand, such as increased stacking between aptamer and phenyl groups on the ligand which holds the ligand more tightly to the aptamer. By contrast, a change in k_on_ is more consistent with a more readily accessible binding pocket for ligand entry. The values measured for k_on_ of both compounds are ~10^4^ to 10^5^ lower than the diffusion-controlled limit of 10^2^ to 10^4^ μM^−1^s^−1^ (Alberty and Hammes 1958). A previously measured k_on_ for DFHBI of 6.2 ± 0.1 × 10^−2^ μM^−1^s^−1^ binding to surface-attached Spinach (Han et al. 2013) is about 300-fold higher than our measurements of DFHBI binding to Spinach2 in solution. However, for both aptamers the k_on_ is orders of magnitude lower than the diffusion-controlled limit. This suggests a large energy barrier to the entry of these compounds into the Spinach2 pocket with the additional pentafluorophenyl group of PFP-DFHBI decreasing that barrier by 10-20 fold. The increased rate of binding of PFP-DFHBI compared with DFHBI may be due to the greater disruption in the region around the binding site caused by PFP-DFHBI compared with DFHBI that was detected by SHAPE analysis.

As a broad range of new roles are attributed to RNAs in cells, opportunities will open for regulating cellular activity with small molecule inhibitors. The findings here with the Spinach2 and Broccoli aptamers suggest k_on_ is the critical kinetic parameter and that the inclusion of poly-fluorophenyl moieties might disrupt a local region of the RNA target at a location such as a bulge or loop. Such disturbance might allow the RNA to overcome a local energy barrier to folding with a resulting alteration in structure and potentially function. These results suggest that the inclusion of poly-fluorophenyl groups on RNA-targeting drugs should be explored for the possibility of increasing drug effectiveness.

## MATERIALS AND METHODS

### Reagents

Buffer IC was formulated to approximate intracellular pH and ionic concentrations based on literature reports for these values (Rorsman et al. 1982; Ammann et al. 1995; Grubbs 2002; Romani 2007; Romani 2013) and contained 13.5 mM NaCl, 150 mM KCl, 0.22 mM Na_2_HPO_4_, 0.44 mM KH_2_PO_4_,120 μM MgCl_2_,120 nM CaCl_2_, 100 μM MgSO_4_, 20 mM HEPES, pH 7.4. Buffer SX, which was used for the Spinach2 aptamer selection (Strack et al. 2013), contained and 125 mM KCl, 5mM MgCl_2,_ 40 mM HEPES, pH 7.4. Buffer SB, which was the buffer for Broccoli selection (Filonov et al. 2014), contained 100 mM KCl, 1 mM MgCl_2_, 40 mM HEPES, pH 7.4. All buffers were prepared with deionized distilled water and the pHs were measured at 24-25°C. Due to the need to dissolve reagents in DMSO, all reaction mixes also contained 5% DMSO. Buffer formulations are shown in Table S2. All inorganic chemicals were from Fisher Scientific.

### Preparation of RNAs

The SPN2A RNA used in this study contains the sequence of Spinach2 (Strack et al. 2013) with the addition of two Gs at the 5’ end that are added by T7 polymerase at transcriptional initiation. Sequences of the RNA aptamers and their variants used in this study are shown in Table S2. RNAs were prepared by *in vitro* transcription with the AmpliScribe™ T7 Flash™ or T7 High Yield *in vitro* transcription kits (Epicenter) from templates created by oligonucleotide annealing and PCR amplification. Alternatively the RNA was prepared by lab made T7 polymerase. The BRC1A aptamer and the chemically modified portions of the SPN2A aptamer were synthesized chemically by Integrated DNA Technologies (IDT, Coralville, IA). These are identified in Table S2.

For SPN2A RNAs with either a 2-aminopurine (2AP) or no purine at position 71, a split construction was used. The two halves of the split SPN2A (sequences specified in Table S2) were those used to create the SPN2A RNA in an equivalent way to which the SPN2A RNA was prepared to determine its structure by X-ray crystallography (Warner et al. 2014). One half of each RNA containing either a substitution of 2AP for A21 or with an apurinic pentose sugar at position 21 (Apu) were chemically synthesized by IDT (Coralville, IA) and maintained at −20°C in autoclaved deionized distilled water (ddH2O) until use. Position A21 in this first half-molecule corresponds to A71 in the full-length SPN2A (Table S2). The second RNA molecule half was synthesized as described for the full-length RNA molecule. The two halves were combined in SX buffer, heated to 95°C for 5 min then slowly cooled to 25°C over a period of one hour. These reassembled aptamers are referred to as split SPN2A (spSPN2A).

### Synthesis of imidazole derivatives

The general procedure for the preparation of chemicals in the small molecule library of DFHBI analogs with alternate imido-N extensions was: To the azlactone (0.200 g, 0.711 mmol) in ethanol (10 mL) was added a solution of the primary amine (0.853 mmol) in ethanol (10 mL) followed by potassium carbonate (0.1474 g, 1.07 mmol). The reaction mixture was refluxed for 12 h. After cooling to room temperature, the solvent was removed by evaporation. Water (15 mL) was added and the pH was adjusted to 3 using 1 M HCl. The solution was left overnight at 4 °C and the resulting precipitated product was captured by filtration. In some cases additional purification by preparative TLC was required. The products were yellow solids. Each compound resolved as one spot on TLC. NMR and high resolution mass spectrometry data were consistent with the assigned structures (supplementary material).

Quantum yields (QY) of DFHBI and PFP-DFHBI were determined in complex with tandem SPN2A relative to acridine orange in SX buffer with 5% DMSO. The results were analyzed using ORIGIN 7 (v7.0552 (B552), www.OriginLab.com) for absorbance deconvolution and the formula below to determine QY:

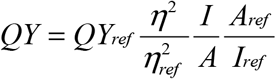

where η = refractive index, A = absorbance, I = integrated area.

### Computational analyses of structure of the aptamer and potential ligands

Images of the Spinach2 structure in association with its ligand were prepared from 4ts2.pdb using Pymol (https://pymol.org/2/). The MM2 force field method, available in the ChemBio3D® Ultra 12 Suite, was used for calculating the properties of organic molecular models from which the 3D images of DFHBI extension ligands were created.

### Fluorescence measurements

Fluorescence intensities were measured by a Cary Eclipse spectrofluorometer (Varian, Palo Alto, CA) for single time point measurements and a Synergy 2 plate reader (Biotek, Winooski, VT) for multiple time point measurements.

#### Determination of affinities

The dissociation constants (K_a_) of the aptamer-ligand complexes were determined by measuring the increase in fluorescence with increasing ligand concentration while the aptamer concentration was held constant. Spinach SPN2A RNA (0.05 μM) was incubated for various time periods (10 min to 3 h) in the dark at 23°C with various concentrations up to 50 μM DFHBI in Buffer SX with 5% DMSO. Broccoli BRC1A RNA (0.05 μM) was incubated under the same conditions with Buffer SB in place of Buffer SX. Fluorescence intensities were measured with λ_ex_ = 485 nm and λ_em_ = 528 nm with 20 nm slit widths. The fluorescence data, normalized to the maximum value in each dataset, was fit in Microsoft Excel using the Solver Add-in (GRG Non-linear method) and the equation F = Fmin+(Fmax*L^n^)/(L^n^+ K_d_^n^) (Huang et al. 2014), where F is fluorescence, L is the concentration of ligand, n is the Hill coefficient, and K_d_ is the binding constant. The fit was calculated by instructing the Solver to minimize the sum of the squares while allowing Fmin, Fmax, n and K_d_ to vary. Ka was calculated as the inverse of the K_d_. For some of the variant aptamers, for which affinity was low, n was set to 1 to obtain a proper fit. Samples of titration data are shown in Fig. S1

### Kinetic measurements

The k_on_ and k_off_ rates for 97SPN2A with DFHBI or PFP-DFHBI were done by stopped-flow kinetics using a MOS-250 spectrometer linked to a stopped-flow SFM-400 apparatus by a MPS-60 microprocessor unit (Bio-Logic, Claix, France) or using a Cary eclipse fluorescence spectrophotometer. The interaction with either ligand was monitored by fluorescence (excitation at 460nm and emission at 500nm). For k_on_ measurements, 20nM 97SPN2A was mixed with DFHBI or PFP-DFHBI in 125 mM KCl, 5 mM MgCl_2_, 40 mM HEPES, 5% DMSO, pH 7.4 at 25_C at final concentrations of 50, 100, 200, 250, 300, 400, 500 or 800 nM. Binding was monitored by the change in the fluorescence at 500 nm over the period from 20 msec to 150 sec after mixing. In some experiments the final concentration of DMSO was 2.5%. For k_off_ rates, GK117 was present in excess to replace the dissociated ligand in the SPN2A binding pocket and thereby prevent the reassociation of DFHBI or PFP-DFHBI. Mixtures of 97SPN2A and DFHBI or PFP-DFHBI were mixed with GK117 to achieve final concentrations at t=0 of 40 mM HEPES, 125 mM KCl, 5 mM MgCl_2_, pH 7.4, 5% DMSO, 100 nM 97SPN2A, 20 μM DFHBI or PFP-DFHBI and GK117 at 100 or 200 μM. The dissociation of DFHBI or PFP-DFHBI from 97SPN2A was monitored by the change in fluorescence at 500 nm during the period of 20 msec to 150 sec. The kinetic data were fit with an exponential function using Costat (version 6.541) to obtain k_off_ and k_obs_. k_on_ is the slope of the linear transformation k_obs_ = k_on_*[L] + c where [L] is the ligand (DFHBI or PFP-DFHBI) concentration and c is a constant.

### SHAPE

SPN2A RNA in water was heated to 65°C for 5 min, cooled to room temperature then diluted into Buffer SX at 37°C to 9.9 μM RNA, with and without 10 μM ligand. The mixtures were incubated at 65^°^C for 10 min. The RNA was reacted with or without 5 mM benzoyl cyanide in Buffer SX with 4.9% DMSO, Reverse transcription was done with Thermo Fisher Superscript IV with 0.5 mM of each nucleotide triphosphate (NTP) and a 5′ hexachloro-fluorescein (HEX) end-labeled primer (TTTTGTTTATTCTTTT). A G-ladder was created using the same primer 5’ labeled with 6-carboxyfluorescein (6FAM) and RT reactions that included 0.5 mM ddGTP. Completed reactions were ethanol precipitated and re-suspended in deionized formamide (Hi-Di formamide, Thermo Fisher) with the G-ladders. Fragmentation analysis of the resulting cDNAs was performed on an Applied Biosystems 3730 DNA Analyzer by the DNA facility (Iowa State University). The data was normalized using the QuShape V 1.0 software with the following calculation:

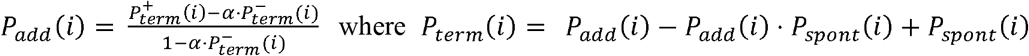

Where 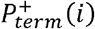 and 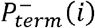 are the probabilities of primer termination at nucleotide i in reaction mixes with (+) and without (−) benzoyl cyanide. The parameter α accounts for scaling differences between the spontaneous termination probabilities in the two reactions (Karabiber et al. 2013).

## SUPPLEMENTAL MATERIAL

Supplemental material is available for this article.

## ACKNOWLEDGEMENTS

We thank Jonathan Beasley for synthesizing DFHBI, Joshua Alterman for preparing a batch of GK117, Aleksei Ananin for help with MS measurements, Scott Nelson for help with stopped-flow measurements, Richard Honzatko for useful discussion and Walter Moss for critiquing an early version of the manuscript. Financial support for this study was provided by grant R21AI114283 from the National Institutes of Health (NIAID) (chemistry) and Aptalogic Inc. (Molecular biology). MNH owns Aptalogic Inc.

## Author Contributions

The chemical library was prepared by IMG with oversight from GAK. The majority of the data was collected and individually analyzed by SMA with contributions from MI and LB. MNH compiled and evaluated the data, developed the hypothesis and directed the work progress. LB discovered the change in aptamer affinity with time, MI proposed the gateway A as important, and MNH proposed the G67 OH group. MNH wrote the manuscript with contributions from all coauthors.

